# The role of embryonic origins and suture proximity in repair of cranial bone

**DOI:** 10.1101/2023.06.27.546687

**Authors:** Daniel Doro, Annie Liu, Jia Shang Lau, Arun Kumar Rajendran, Christopher Healy, Agamemnon E. Grigoriadis, Sachiko Iseki, Karen J. Liu

## Abstract

The cranial sutures are proposed to be a stem cell niche, harbouring skeletal stem cells (SSCs) that are directly involved in development, homeostasis, and healing. Like the craniofacial bones, the sutures are formed from both mesoderm and neural crest. During cranial repair, neural crest cells have been proposed to be key players; however, neural crest contributions to the sutures are not well defined, and the relative importance of suture proximity is unclear. Here, we combine calvarial wounding experiments with genetically labelled mouse models to demonstrate that suture proximity predicts the efficiency of cranial repair. Furthermore, we define the neural crest-mesoderm boundary within the sutures and demonstrate that *Gli1*+ and *Axin2*+ SSCs are present in all calvarial sutures examined. We propose that the position of the defect determines the availability of neural crest-derived progenitors, which appear to be a key element in the repair of calvarial defects.

**Summary statement:** This study provides a new examination of the embryonic origins of cranial sutures in mouse. Using genetic lineage labelling, we identify specific contributions of neural crest cells and associated skeletal stem cells during repair of calvarial wounds. This is extremely relevant considering recent findings regarding the role of suture-residing stem cells.

## Introduction

Craniofacial bone repair, which can be necessary due to accidents, congenital anomalies, or diseases such as cancer, is a daunting clinical challenge and a significant biomedical burden. Current treatment strategies include replacements; however, these do not truly mimic bone and can lead to difficulties in integration with other tissues such as muscle. Poor healing can have severe impact on function and aesthetics, while multi-fragment breaks can be difficult to reconstruct^1^. Therefore, there is a need to better understand the endogenous craniofacial repair process, including a clearer definition of the osteogenic stem cell niche.

In craniofacial structures, the calvarial and facial bones form via intramembranous ossification, in which osteoblasts coalesce and differentiate directly within a membrane, without a cartilaginous intermediate^2^. Growth of these skull bones is organized at fibrous joints called sutures and occurs at the leading edges of the separate bones^2^. Throughout embryonic and postnatal development, the sutures remain as active sites of bone formation. By early adulthood, when growth of the skull is complete, sutures become quiescent and gradually fuse. Nevertheless, cells residing in the sutures or adjacent periosteum retain some limited potential to heal calvarial bone in adults, following injury or disease^3^.

Here, we make use of a critical and underappreciated observation: that adult frontal bones in the mouse heal more efficiently than parietal bones^4,5^. Studies from mouse models have demonstrated that the embryonic frontal bones are of neural crest origin, while the parietal bones are mesodermally-derived^6^, raising the possibility that developmental history influences osteogenic potential. Indeed, we have demonstrated that neural crest-derived osteoblasts have an increased differentiation capacity^5^. Anecdotal evidence suggests a similarly improved healing capacity in human frontal bones^7^. These observations raise the possibility that there are increased numbers (or activity) of calvarial stem cells in sutures adjacent to the frontal bones.

Most of the original studies comparing frontal and parietal healing efficiency do not account for the role of suture-derived osteoprogenitors, or the relevance of defect position in relation to the sutures. More recently, it has been demonstrated that bone healing is not an evenly distributed event across the parietal bone surface^8^. While these studies show a correlation between repair efficiency and suture proximity, they disregard the dual embryonic origin of the cranial sutures and the possibility that distinct lineage contributions may influence repair. Furthermore, several studies suggest that neural crest-derived cells, and not the mesoderm, play key roles in the pathogenesis of craniofacial malformations^9–11^, but also that neural crest may account for differences in cranial healing efficiency upon injuries of every sort^4,12,13^.

Based on historical lineage tracing assays using *Wnt1::cre*-driven ROSA26-lacZ reporters^6,11,14^, both interfrontal and sagittal sutures are neural crest-derived, while the coronal suture is mesodermal. These studies focused on late embryonic/early post-natal stages, when cranial sutures are not yet clearly defined. To date, no definitive description of the neural crest-mesoderm boundary in the murine cranial vault has been established. Our overall hypothesis is that, in addition to suture proximity, the embryonic origins of the cranial sutures are also relevant when assessing repair. We hypothesise that calvarial sutures, notably those with majority neural crest contributions, may harbour more substantial skeletal stem cell populations. Thus, for non-neural crest derived bone, such as the parietal bones, proximity to specific sutures is crucial for homeostasis as well as cranial repair.

In this study, we first confirm that murine frontal bones heal more efficiently than parietal bones and note that the position of the defect in relation to the sutures is paramount for the outcome. We use genetically labelled lineage markers, to define the neural crest-mesoderm domains of the cranial sutures and the presence of *Axin2+* and *Gli1+* stem cell populations therein. We then demonstrate that *Wnt1::cre+* neural crest lineages are found in every subcritical defect regardless of the embryonic origin of the wounded bone, or the proximity to a neural crest suture. This suggests that localised sources of neural crest cells play a crucial role in healing of mesodermal skull bones.

## Results and Discussion

### Suture proximity determines the outcome of a subcritical cranial defect

While regional differences in repair efficiency have previously been reported, the dura mater and periosteum were thought to be the main sources contributing osteoprogenitors for cranial repair^15,16^. Here we set out to determine the role of proximity to the sutural niche. 1 mm subcritical defects (termed midfrontal and midparietal) were made in the frontal bone of adult mice postnatal day 40 (PN40), equidistant to the interfrontal and coronal sutures, as well as in the parietal bone, equidistant to the lambdoid, squamous, sagittal, and coronal sutures (Figure 1A – empty dotted outlines, 1B). After 4 weeks, we observed a striking difference between midfrontal and midparietal repair (Figure 1D), which accords with previous observations^4,12,13,17^. When the defects are made proximally to the sutures (peri-coronal frontal, peri-coronal parietal and peri-lambdoid) (Figure 1C), no significant difference in repair is seen between frontal and parietal bones (Figure 1D), whereas the peri-coronal parietal defect shows significantly more repair than the midparietal (Figure 1D). Our findings corroborate recent studies, which show that subcritical parietal defects are more likely to heal the closer they are to the cranial sutures^8^. However, we do not see any specific increase in repair in the vicinity of a specific suture in relation to the other. This shows that that even though the sutures may contribute differently for cranial repair, the proximity to any suture is enough to provide subcritical healing, while the more distant parietal defect (here termed midparietal) seems to exceed the minimum critical distance from a suture.

**Figure 1:**
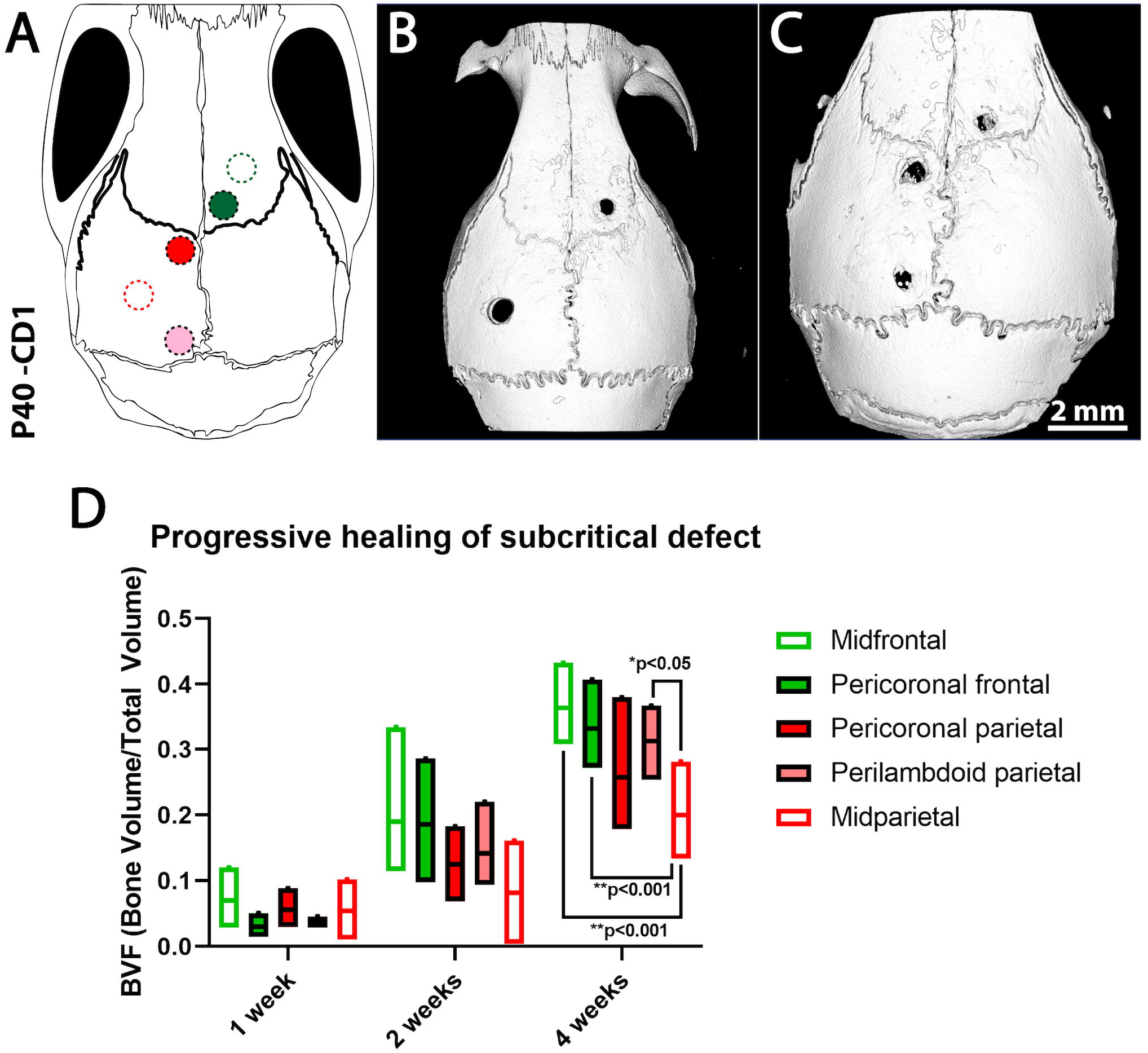
Subcritical repair efficiency correlates with suture proximity irrespective of the calvarial bone. (A) Schematic shows the position of cranial defects in relation to adjacent sutures. Green outline – midfrontal, Solid green – Peri-coronal frontal, Solid red – Peri-coronal parietal, Red outline – midparietal, Solid pink – peri-lambdoid. (B, C) Top view microCT scan of CD1 mouse heads 4 weeks after 1 mm subcritical defect surgery. (B) Defects equidistant from surrounding sutures in right frontal and left parietal bones. (C) Defects proximal to coronal and interfrontal suture (right), coronal and sagittal suture (top left), sagittal and lambdoid suture (bottom left). (D) Bone Volume Fraction (BVF) analysis over 4-week period post-subcritical defect surgery. Two-tailed unpaired t test was used. n=6. *p<0.05, **p<0.001, ***p<0.001.

### Neural crest-mesoderm boundary separates cranial sutures into two domains

In mouse, *Wnt1::*cre-dependent lineage labelling has demonstrated the neural crest-origin of the frontal bones and the *Mesp1::cre-*labelled mesoderm contributing to the parietal bones^6,11,14^. However, the embryonic origins of the cranial sutures are debatable, as they rely on early observations that lacked resolution. Nevertheless, we and others have repeatedly confirmed a higher osteogenic potential of neural crest osteoblasts when compared to mesoderm osteoblasts^4,5,18,19^. We sought to determine the neural crest domain and the presence of stem cell lineages in all calvarial sutures using high resolution-lineage tracing.

Transgenic *Wnt1::cre* mice were crossed with mice carrying a cre-responsive *Rosa26R*^*mTmG*^ reporter. In these mice, *Wnt1::cre* expression begins approximately at embryonic day 8.5 (E8.5) in the dorsal neural tube, just as cranial neural crest cells are induced^27^. Soon after, these cells migrate into the cranial vault and can be tracked with membrane green fluorescent protein (mGFP). Cells that do not express lacking *Wnt1::cre* are labelled with membrane tomato (mTom) and are presumptive mesoderm. As expected, the frontal bone shows exclusive neural crest contribution (mGFP+) as opposed to the parietal bones, which is of mesodermal origin, lacking mGFP (Figure 2A). When we examine the midline sutures, we see a clearly defined neural crest-mesoderm boundary, which lies right at the middle of the sagittal suture, delimiting a neural crest domain (interfrontal, coronal and anterior-sagittal suture) and a mesoderm domain (posterior-sagittal and lambdoid suture) (Figure 2A, 2B). A coronal section at a more anterior part of the sagittal suture shows complete absence of mesoderm (Figure 2B, 2C-E), contrasting a more posterior section which shows very few neural crest cells with abundant mesoderm (Figure 2F-H, I-K). Likewise, the interfrontal suture shows only neural crest (Figure 2C-E), whereas the lambdoid suture shows only mesoderm (Figure 2B, 2L-N). Altogether, this shows that the sagittal suture has dual embryonic origin with a well-defined neural crest-mesoderm boundary that extends past the anterior edge of the parietal bones, recapitulating the pattern observed in early development. This separates the sutures into two domains: The neural crest-derived sutures (interfrontal, coronal, squamous and anterior sagittal) and the mesodermal sutures (lambdoid and posterior sagittal).

**Figure 2:**
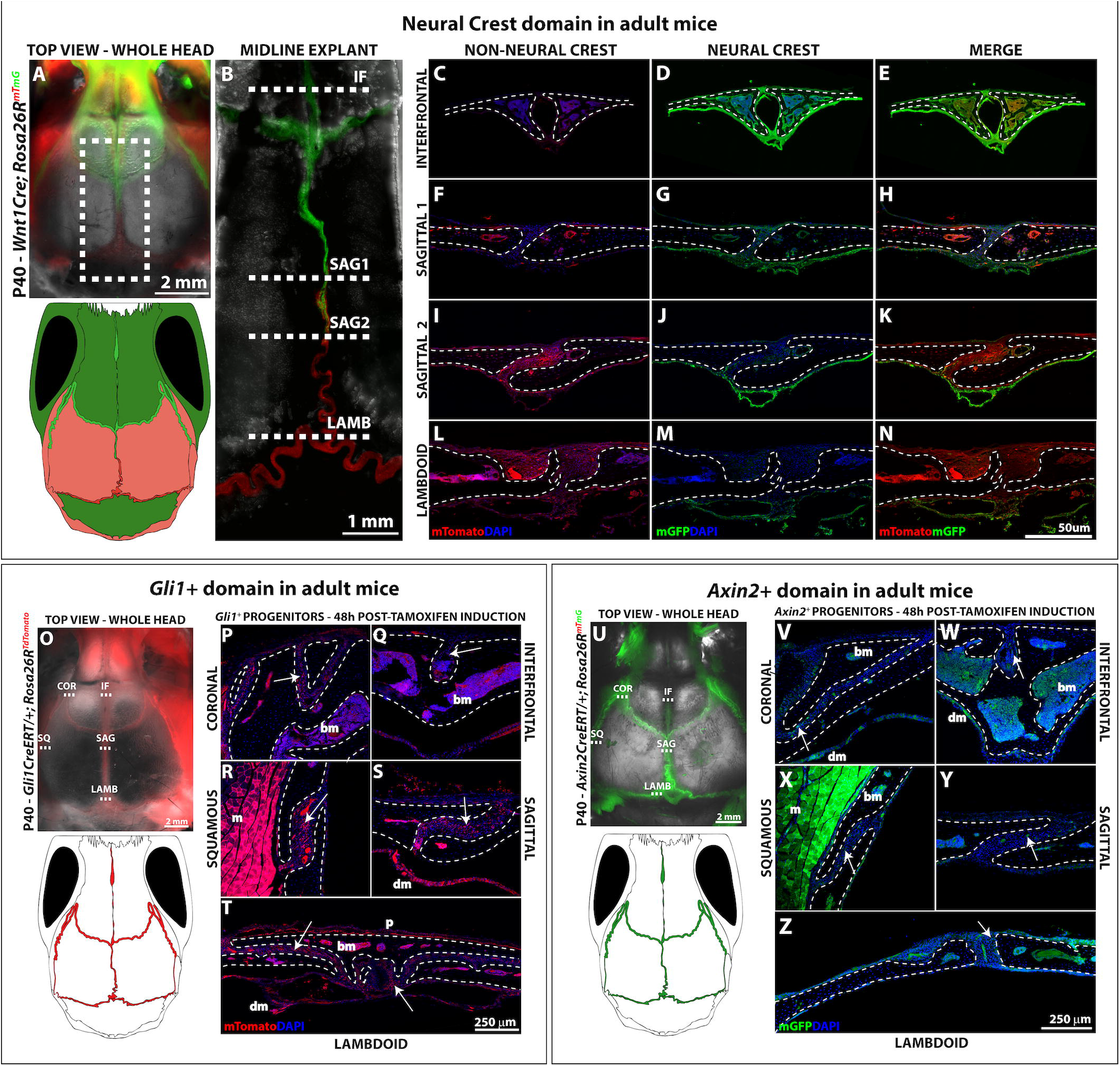
Mapping the identity of cranial sutures. (A) Top view of P40 *Wnt1Cre*; *Rosa26R*^mTmG^mouse. Grey – Brightfield, Green – Neural Crest, Red – Mesoderm. Dashed box shows area of the midline explant in C. Scale bar: 2mm. Schematic depicts the neural crest-mesoderm domain in the cranial vault based on the linage tracing of A. (B) Confocal scan of the midline explant confirming the neural crest-mesoderm boundary at the middle of the sagittal suture. (C-N) Confocal scans of coronal sections at different regions of the midline showing non-neural crest tissue (C,F,I,L – red), neural-crest tissues (D,G,J,M – green) and merged channels. Dashed lines show estimated planes of section in C-N. IF – Interfrontal suture (C-E), SAG1 – Sagittal suture region 1 (F-H), SAG2 – Sagittal suture region 2 (I-K), LAMB – Lambdoid suture (L-N). Scale bar: 1 mm. (E,H,K,N). Nuclear staining is shown in blue. n=3. Scale bar: 50 µm. (O) Top view picture of 40 day-old *Gli1CreERT*/+; *Rosa26R*^TdTomato^ mouse 48 h after tamoxifen induction. Grey – Brightfield, Red – *Gli1+* domain. COR – Coronal suture, IF – Interfrontal suture, SQ – Squamous suture, SAG – Sagittal suture, LAMB – Lambdoid suture. Scale bar: 2 mm; Schematics show *Gli1+* domain in the transgenic mouse calvarium. (P-T) Coronal sections of the calvarial sutures at the regions marked in O. Dashed lines show the bone outline. *Gli1+* are in red. Scale bar: 250 µm. (U) Top view picture of 40 days old *Axin2CreERT*/+; *Rosa26R*^*mTmG*^mouse 48 h after tamoxifen induction. (red channel not shown). Grey – Brightfield, Green – *Axin2+* domain. COR – Coronal suture, IF – Interfrontal suture, SQ – Squamous suture, SAG – Sagittal suture, LAMB – Lambdoid suture. Scale bar: 2 mm; Schematics show *Axin2+* domain in the transgenic mouse calvarium. (V-Z) Coronal sections of the calvarial sutures at the regions marked in U. Dashed lines show the bone outline. *Axin2+* cells are in green. In O-Z white arrowheads show red (*Gli1+*) or green (*Axin2+*) positive cells. bm – bone marrow, m – muscle, dm – dura mater and p -periosteum are notoriously auto fluorescent tissues. Scale bar: 250 µm

### Stem cell populations are present in every calvarial suture and contribute to calvarial repair

Several putative calvarial stem cell markers have been identified: the Hedgehog-pathway transcription factor *Gli1*^20^, the Wnt-responsive gene *Axin2*^21^ and the transcription factor *Prx1*^22^. Here, we investigate the presence of *Gli1* and *Axin2* stem cell populations using *Gli1::creER*^*T2*^ *or Axin2::creER* drivers in combination with *Rosa26*^Tomato^ and *Rosa26R*^mTmG^ reporters, respectively. To label *Gli1* and *Axin2* populations 38 day-old mice were given tamoxifen and the heads were collected 2 days after injection. Top view and coronal sections of *Gli1*-*CreER*^*T2*^; *Rosa26*^Tomato/+^ heads reveal the abundant presence of *Gli1+* cells in all calvarial sutures (Figure 2O-T). The analogous experiment with *Axin2-CreER*^*T2*^; *Rosa26R*^+/mTmG^ animals shows *Axin2+* cells also detected in every suture; however, they are sparse in comparison to *Gli1+* (compare Figure 2U-Z to 2O-T), consistent with prior suggestions that the *Axin2+* cells could be a subset of *Gli1+* progenitors ^21,22^. Overall, we see no apparent difference in *Gli1+* and *Axin2+* cell content in the cranial sutures examined.

To investigate whether *Gli1+* osteoprogenitors could populate all healing defects, subcritical defects were made at 5 sites of *Gli1*-*CreER*^*T2*^; *Rosa26*^Tomato/+^ mice as indicated (Figure 3A) and *Gli1+* cells were examined in healing wounds 1 week post-surgery. We found that all wounds had *Gli1+* cells, regardless of the proximity to the sutures. This suggests that although midparietal defects are substantially more distant to the sutures than the other defects, they are still supplied with suture-derived osteoprogenitors, revealing the reparative capability of the suture mesenchyme in relation to distant defects. Importantly, a quantitative assessment of the cell content in these wounds might reveal a correlation of distance vs. cell contribution.

**Figure 3:**
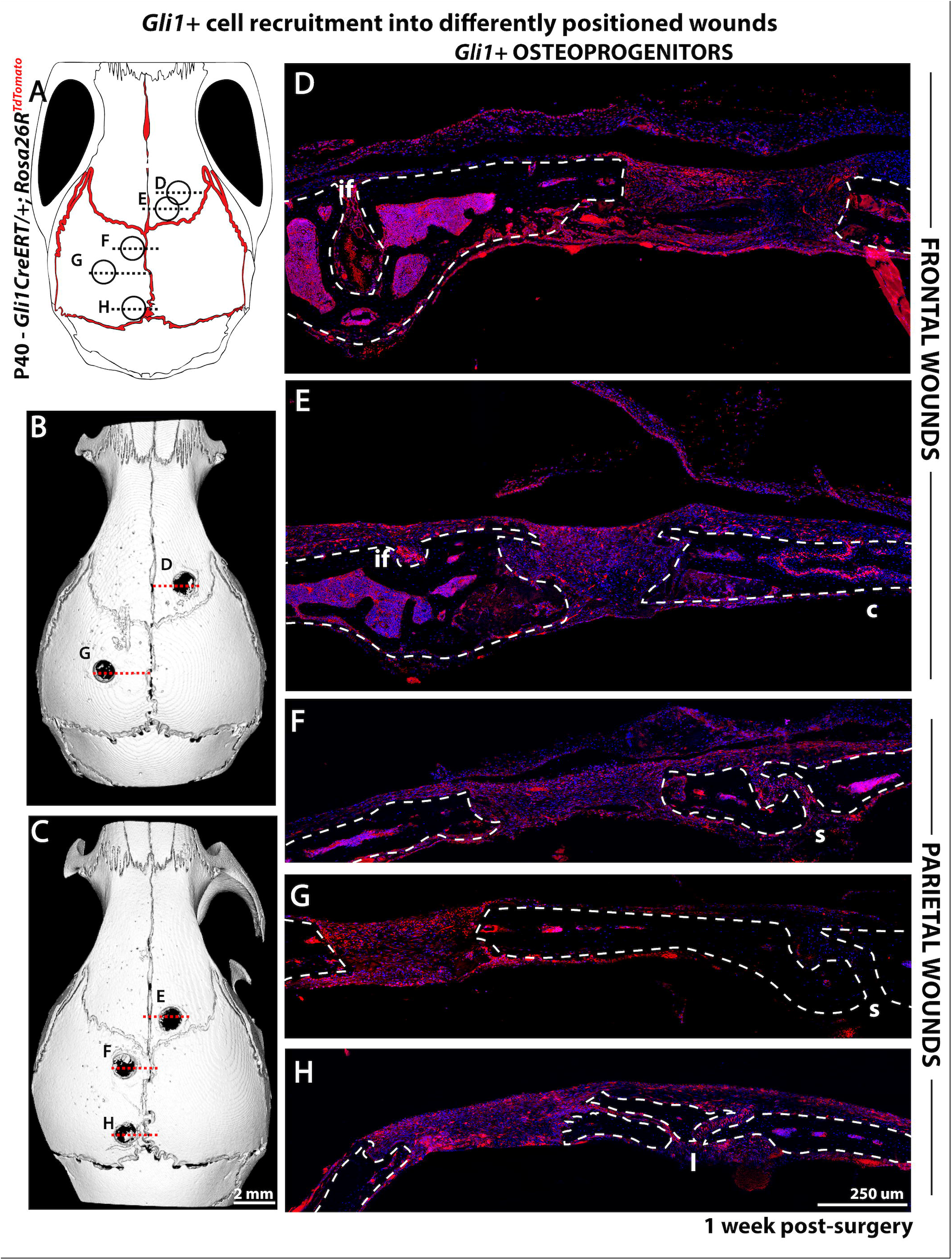
*Gli1+* cells are found in every subcritical defect after 1 week. (A) Schematics show sites of 1 mm subcritical defects in P40 *Gli1CreERT*/+; *Rosa26R*^TdTomato^mice. Circular outlines mark the positions of the wounds in relation to the different bones and sutures. (B, C) Top view CT scans of heads after 1 week from the surgical procedure. Red dashed lines correspond to the predicted plane of sections in D-H. (D-H) Confocal scans of coronal sections at different wound sites 1-week post-surgery. Red – *Gli1+* cells. Dashed lines – Bone outlines. if – Interfrontal suture, c-Coronal suture, s – Sagittal suture, l – Lambdoid suture. Scale bar: 250 µm

### Neural crest contributes progenitors to any cranial defect

Finally, we assessed the extent to which *Wnt1::cre-*expressing neural crest cells infiltrate the repair site after injury. Subcritical defects were made at 6 distinct locations in *Wnt1::cre; Rosa26R*^+/mTmG^ mice (Figure 4A). Defects varied from midfrontal (G) and coronal-frontal (H), which are surrounded by neural crest-derived interfrontal and coronal sutures, to lambdoid-parietal (L) which is only surrounded by mesoderm-derived sutures. 1 week after craniotomy, *Wnt1::cre+* cells were detected around and/or within all defects (Figure 4B-F). This is clearly observed in coronal sections (Figure 4G-I). Even the defects in bones that are mesodermal in origin are filled with a large proportion of mGFP+ cells, suggesting the infiltration of neural crest-derived progenitors (Figure 4G-L). Importantly, within the frontal bone outlines, every cell is expected to be GFP+ (white arrowheads) as these bones are of neural crest origin (Figure 4g’, h’). In contrast, within parietal bone outlines we observe a mix of GFP-(yellow arrowheads) and GFP+ (white arrowheads) (Figure 4i’, j’, l’) suggesting that new bone was formed from neural crest-derived progenitors, since parietal bones are expected to be mesodermal (GFP-). One exception is the midparietal defect (Figure 3k’), which has maintained the original flat edges, as obtained by the cylindrical nature of the drill bit, indicating minimal repair. This wound has a complete absence of green cells within the bone outline. This is consistent with previous observations of poor healing capacity of the mid-parietal bone region.

**Figure 4:**
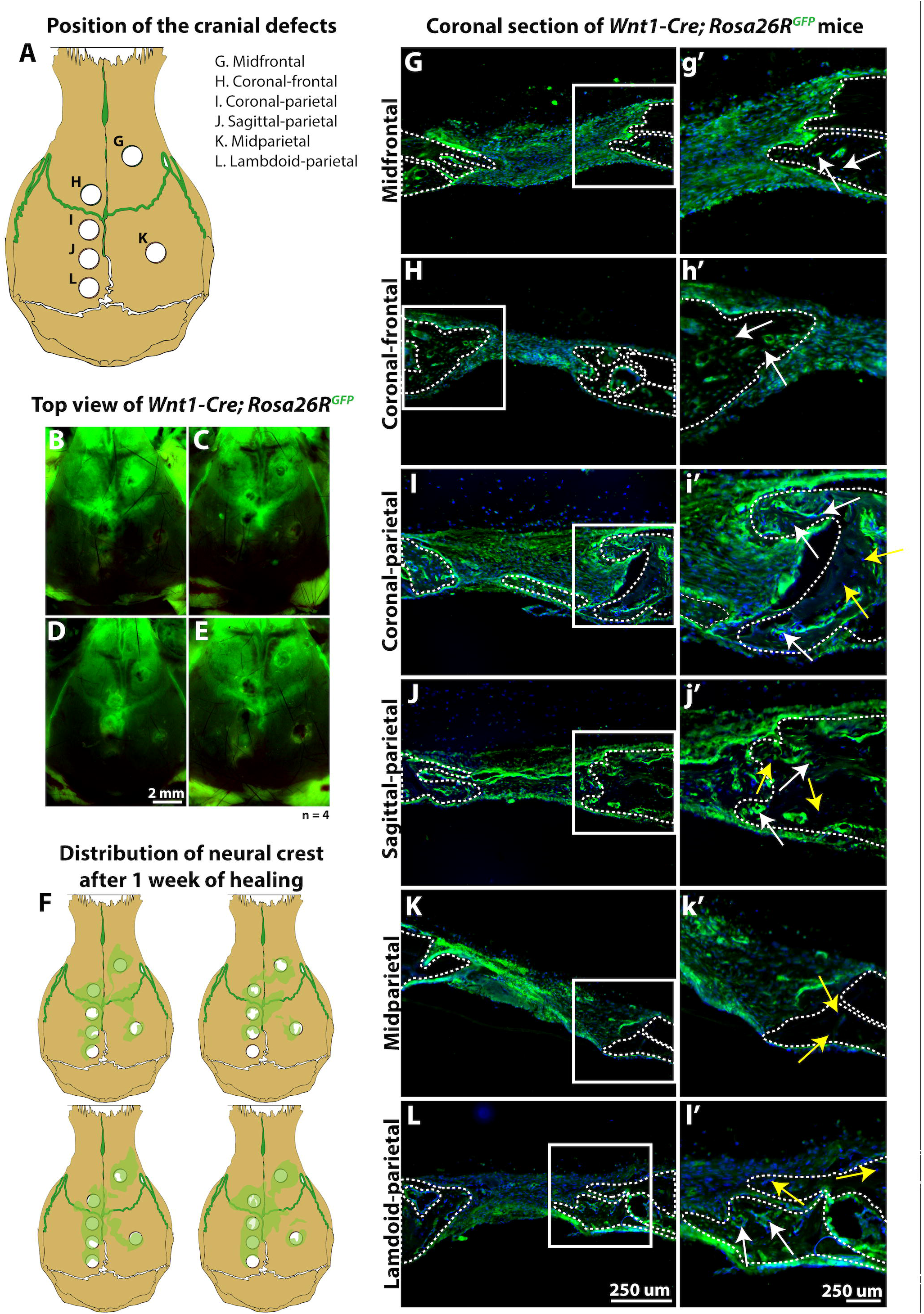
Neural crest cells are recruited to any cranial subcritical defect irrespective of the embryonic origins of the bone or surrounding sutures. (A) Schematics showing the position of 6 subcritical defects (1 mm). (B-E) Top view of 40 days-old *Wnt1-Cre; Rosa26R*^*GFP*^ mice 1 week after the surgical procedure. Green -neural crest derived tissue. (F) Schematics show the distribution of neural crest cells across the calvarial surface based on the fluorescence observed in B-E. (G-L) Coronal sections at the centre of each defect described in A. Neural crest-derived cells are shown in green, nuclear staining in blue. Dashed lines outline the bones, and the boxed area is shown in high magnification on panels g’-l’. Yellow arrowheads -non-neural crest-derived cells, White arrowheads -neural crest-derived cells.

While the osteogenic potential of neural crest versus mesoderm sutures has not yet been defined, it is intriguing that *Wnt1::cre*+ neural crest cells seem to contribute to the repair of every defect, including those in a mesoderm-dominant area (Figure 4). While we propose that the suture proximity is a key factor in repair capacity, it is also worth noting that the underlying dura mater is also neural crest derived, and a recent study has shown contributions of dura mater cells to suture regeneration^23^. Regardless, neural crest-derived cells appear to be required for efficient repair, supporting previous observations that neural crest osteoprogenitors have higher osteogenic capabilities than other mesodermally-derived calvarial cell populations.

In conclusion, we have revisited the implications of embryonic identity and spatial positioning of cranial defects in relation to sutures. While the embryonic origins of the cranial bone appear to predict of the outcome of defect repair, it is important to consider the neural crest composition of the intervening sutures, and the presence of suture-residing osteoprogenitors relative to the calvarial wound site.

## Materials & Methods

### Animal Procedures

All procedures were approved by King’s College London ethical review process and performed in accordance with UK Home Office guidelines Project Licence P8D5E2773 (KJL) or by the Institutional Animal Care and Use Committee of Tokyo Medical and Dental University (A2019-060C3 (SI)).

Mouse lines: CD-1 mice were obtained from Charles River Laboratories. *Rosa26R*^*mTmG*^, *Rosa26R*^*TdTomato*^, *Rosa26R-eGFP* mouse reporter lines have all been described previously^24–26^. The following Cre drivers were used: *Wnt1cre, Axin2*^*creERT*^, *Gli1cre*^*ERT2* 27–29^.

### Subcritical defects

Postnatal day 40 mice were weighed and anesthetized with appropriate dose (10^μ^L/g of body weight) of the 10mg/mL Ketamine/2mg/mL Xylazine cocktail. (Vetalar®/Rompun®). The state of deep anaesthesia was confirmed through tail flick test and hind paw withdrawal response. Once the animals were heavily sedated, the fur on the top of the head was shaved using a hair trimmer. A sagittal incision was performed along the midline with a scalpel. The periosteal layer was then removed with the help of a cotton bud and 1 mm defects were drilled at top speed (50.000RPM) into frontal and parietal bones of the mouse skull using a dental hand drill (Handy-ECO 1000 – Marathon®). The defects were very carefully performed to avoid injuring the dura mater. The bone surface was then rinsed with sterile PBS to remove any debris and the sagittal incision was sutured with 6-0 ETHILON® Nylon absorbable suture. Finally, the mice were moved to a 28°C incubator until full recovery.

### Tamoxifen injection

Targeted deletion in the Cre^ERT^ mice was performed by peritoneal injection of a 10mg/mL Tamoxifen solution (Sigma®) in the adult mouse at the desired stage. The solution was previously prepared by dissolving 10mg of Tamoxifen into 100^μ^L of absolute ethanol and 900^μ^L of corn oil. The dosage was determined according to the following: 1.5mM/g of body weight. Volume: 7,5x g of body weight (^μ^L)).

### Micro-CT scanning

All head samples were fixed for 48 h at room temperature in 4% PFA and scanned using a Scanco Medical µCT50® with the following settings: Energy 70 kV, Intensity 114 µA.

### Sample fixation and sectioning

Samples were fixed in 4% paraformaldehyde (PFA) for 48 hours at 4°C. After 3 PBS washes, the skull cap was dissected and decalcified in 10% formic acid. After 3 more PBS washes, the samples were moved to a 30% sucrose solution in PBS until they sunk. The embedding solution was then replaced with a 30% sucrose solution mixed (1:1) with OCT compound (CellPath®). The samples were incubated at 4°C for another 48 hours. Finally, the samples were moved and oriented in a plastic Tissue-Tek® Cryomold® filled with OCT compound. The mould was quickly moved into a dry ice bath with absolute ethanol until the OCT block was fully solidified. Cryosections were performed using OFT5000® cryostat microtome (Bright Instruments®). 15μm sections were mounted into Superfrost Ultra Plus® slides (ThermoScientific®), which were then stored at room temperature for future use.

### Microscopy

A stereoscope (Nikon SMZ1500) with an attached camera (Nikon digital sight DS-Fi1) was used to take top view fluorescent pictures of whole heads.

Confocal microscopy was performed on a Leica Microsystems CMS GmbH TCS SP5 DM16000. Image sequences were reconstructed using FIJI (Image J) analysis software.

